# Structural analysis of SARS-CoV-2 genome and predictions of the human interactome

**DOI:** 10.1101/2020.03.28.013789

**Authors:** Andrea Vandelli, Michele Monti, Edoardo Milanetti, Alexandros Armaos, Jakob Rupert, Elsa Zacco, Elias Bechara, Riccardo Delli Ponti, Gian Gaetano Tartaglia

**Author notes:** to whom correspondence should be addressed to (RDP) and or (GGT).

## Abstract

Specific elements of viral genomes regulate interactions within host cells. Here, we calculated the secondary structure content of >2000 coronaviruses and computed >100000 human protein interactions with severe acute respiratory syndrome coronavirus 2 (SARS-CoV-2). The genomic regions display different degrees of conservation. SARS-CoV-2 domain encompassing nucleotides 22500 – 23000 is conserved both at the sequence and structural level. The regions upstream and downstream, however, vary significantly. This part codes for the Spike S protein that interacts with the human receptor angiotensin-converting enzyme 2 (ACE2). Thus, variability of Spike S may be connected to different levels of viral entry in human cells within the population.

Our predictions indicate that the 5’ end of SARS-CoV-2 is highly structured and interacts with several human proteins. The binding proteins are involved in viral RNA processing such as double-stranded RNA specific editases and ATP-dependent RNA-helicases and have strong propensity to form stress granules and phase-separated complexes. We propose that these proteins, also implicated in viral infections such as HIV, are selectively recruited by SARS-CoV-2 genome to alter transcriptional and post-transcriptional regulation of host cells and to promote viral replication.

## INTRODUCTION

A disease named Covid-19 by the World Health Organization and caused by the severe acute respiratory syndrome coronavirus 2 (SARS-CoV-2) has been recognized as responsible for the pneumonia outbreak that started in December, 2019 in Wuhan City, Hubei, China ^1^ and spread in February to Milan, Lombardy, Italy ^2^ becoming pandemic. As of June 2020, the virus infected >9’000’000 people in >300 countries.

SARS-CoV-2 is a positive-sense single-stranded RNA virus that shares similarities with other beta-coronavirus such as severe acute respiratory syndrome coronavirus (SARS-CoV) and Middle East respiratory syndrome coronavirus (MERS-CoV) ^3^. Bats have been identified as the primary host for SARS-CoV and SARS-CoV-2 ^4,5^ but the intermediate host linking SARS-CoV-2 to humans is still unknown, although a recent report indicates that pangolins could be involved ^6^.

Coronaviruses use species-specific proteins to mediate the entry in the host cell and the spike S protein activates the infection in human respiratory epithelial cells in SARS-CoV, MERS-CoV and SARS-CoV-2 ^7^. Spike S is assembled as a trimer and contains around 1,300 amino acids within each unit ^8,9^. The receptor binding domain (RBD) of Spike S, which contains around 300 amino acids, mediates the binding with angiotensin-converting enzyme, (ACE2) attacking respiratory cells. Another region upstream of the RBD, present in MERS-CoV but not in SARS-CoV, is involved in the adhesion to sialic acid-containing oligosaccharides and plays a key role in regulating viral infection ^7,10^.

At present, few molecular details are available on SARS-CoV-2 and its interactions with the human host, which are mediated by specific RNA elements ^11^. To study the RNA structural content, we used *CROSS* ^12^ that was previously developed to investigate large transcripts such as the human immunodeficiency virus HIV-1 ^13^. *CROSS* predicts the structural profile of RNA molecules (single-and double-stranded state) at single-nucleotide resolution using sequence information only. Here, we performed sequence and structural alignments among >60 SARS-CoV-2 strains and identified the conservation of specific elements in the spike S region, which provides clues on the evolution of domains involved in the binding to ACE2 and sialic acid.

As highly structured regions of RNA molecules have strong propensity to form stable contacts with proteins ^14^ and promote assembly of specific complexes ^15,16^, SARS-CoV-2 domains enriched in double-stranded content are expected to establish interactions within host cells that are important to replicate the virus ^17^. To investigate the interactions of SARS-CoV-2 RNA with human proteins, we employed *cat*RAPID ^18,19^. *cat*RAPID ^20^ estimates the binding potential of a specific protein for an RNA molecule through van der Waals, hydrogen bonding and secondary structure propensities allowing identification of interaction partners with high confidence ^21^. The computational analysis of more than 100000 interactions with SARS-CoV-2 RNA reveals that the 5’ end of SARS-CoV-2 has strong propensity to bind to human proteins involved in viral infection and reported to be associated with HIV infection. A comparison between SARS-CoV and HIV reveals similarities ^22^ that are is still unexplored. Interestingly, HIV and SARS-CoV-2, but not SARS-CoV nor MERS-CoV, have a furin-cleavage site occurring in the spike S protein, which could explain the high velocity spread of SARS-CoV-2 compared to SARS-CoV and MERS-CoV ^23,24^.

Many processes related to SARS-CoV-2 replication are at present unknown and our study aims to suggest relevant protein interactions for further investigation. We hope that our large-scale calculations of structural properties and binding partners of SARS-CoV-2 will be useful to identify the mechanisms of virus replication within the human host.

## RESULTS

### SARS-CoV-2 contains highly structured elements

Structured elements within RNA molecules attract proteins ^14^ and reveal regions important for interactions with the host ^25^. Indeed, each gene expressed from SARS-CoV-2 is preceded by conserved transcription-regulating sequences that act as signal for the transcription complex during the synthesis of the RNA minus strand to promote a strand transfer to the leader region to resume the synthesis. This process is named discontinuous extension of the minus strand and is a variant of similarity-assisted template switching that operates during viral RNA recombination ^17^.

To analyze SARS-CoV-2 structure (reference Wuhan strain MN908947.3), we employed *CROSS* ^12^ to predict the double-and single-stranded content of RNA genomes such as HIV-1 ^13^. We found the highest density of double-stranded regions in the 5’ end (nucleotides 1-253), membrane M protein (nucleotides 26523-27191), spike S protein (nucleotides 21563-25384), and nucleocapsid N protein (nucleotides 28274-29533; **Fig. 1A**) ^26^. The lowest density of double-stranded regions were observed at nucleotides 6000-6250 and 20000-21500 and correspond to the regions between the non-structural proteins nsp14 and nsp15 and the upstream region of the spike surface protein S (**Fig. 1**) ^26^. In addition to the maximum corresponding to nucleotides 22500-23000, the structural content of Spike S protein shows minima at around nucleotides 21500-22000 and 23500-24000 (**Fig. 1**). We used the *Vienna* method ^27^ to further investigate the RNA secondary structure of specific regions identified with *CROSS* ^13^. Employing a 100-nucleotide window centered around *CROSS* maxima and minima, we found good match between *CROSS* scores and Vienna free energies (**Fig. 1**).

**Fig. 1.**
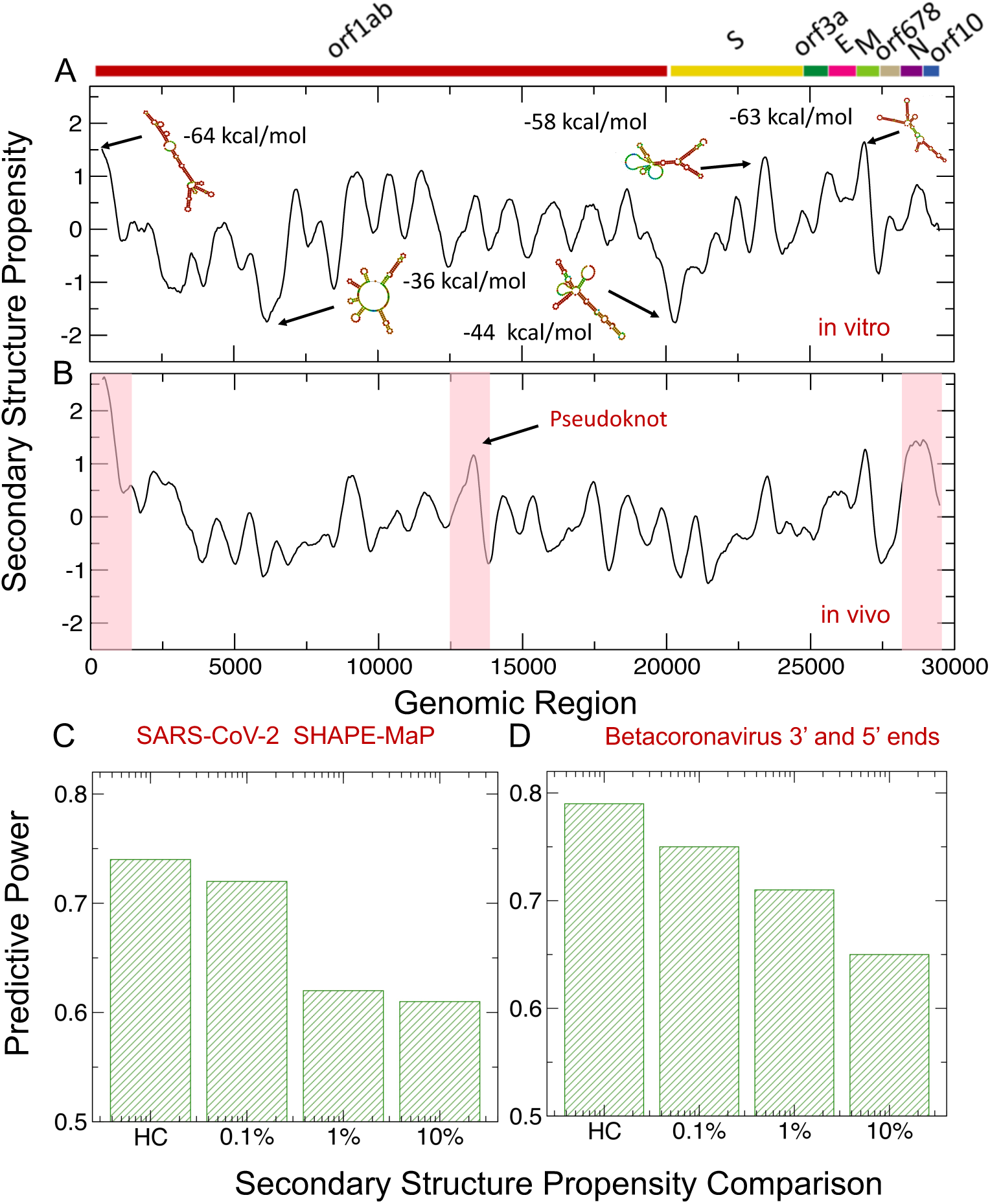
Predictions of SARS-CoV-2 structure. (**A**) Using the CROSS approach ^13,12^, (**A**) we predicted the structural content of SARS-CoV-2 in vitro. We found the highest density of double-stranded regions in the 5’ end (nucleotides 1-250) and within membrane M protein (nucleotides 26500-27000), and spike S protein (nucleotides 22500-23000 regions. Regions with the highest structural content are predicted by Vienna to have the lowest free energies. (**B**) Using CROSS alive ^28^, we studied the structural content of SARS-CoV-2 in vivo. The 5’ and 3’ ends (indicated by red boxes) are predicted to be highly structured. In addition, nucleotides 22500-23000 in Spike S region and nucleotides 13400-13600 (indicated by a red box) forming a pseudoknot ^29^ show high density of contacts. (**C**) Comparison of CROSS predictions with the secondary structure landscape of SARS-CoV-2 revealed by SHAPEMaP ^32^. From low (10%) to high (0.1%) confidence scores, the predictive power, measured as the Area Under the Curve (AUC) of Receiver Operating Characteristics (ROC), increases monotonically (HC corresponds to 10 nucleotides with highest/lowest scores). (**D**) CROSS performances on betacoronavirus 5’ and 3’ ends ^33–36^. Using different confidence scores, we show that CROSS is able to identify double and single stranded regions with great predictive power.

RNA structure *in vitro* and *in vivo* could be significantly different due to the interaction with proteins and other molecules ^28^. Using *CROSS alive* to predict the double-and single-stranded content of SARS-CoV-2 in the cellular context ^28^, we found that both the 5’ and 3’ ends are the most structured regions followed by nucleotides 22500-23000 in the Spike S region, while nucleotides 6000-6250 and 20000-21500 have the lowest density of double-stranded regions (**Fig. 1B**). The region corresponding to nucleotides 13400-13600 shows high density of contacts. This part of SARS-CoV-2 sequence has been proposed to form a pseudoknot ^29^ that is also visible in *CROSS* profile (**Fig. 1A**), but *CROSS alive* is able to identify long range interactions and better identifies the region. Additionally, we used the *RF-Fold* algorithm of the *RNAFramework* suite ^30^ (**Materials and Methods**) to search for pseudoknots. Employing *CROSS* as a soft-constraint for RF-Fold, we predicted 6 pseudoknots (nucleotides 3394-3404, 13723-13732, 14677-14711, 16867-16905, 24844-24884, 27969-27990). The pseudoknot at nucleotides 13723-13732 is in close proximity to the one proposed for SARS-CoV-2 ^29^ and the one at nucleotides 27969-27990 is at the 3’ end, where pseudonoknots have been shown to occur in coronaviruses ^31^.

To validate our results, we compared *CROSS* predictions of double-and single-stranded content predicted (as released in March 2020) with the secondary structure landscape of SARS-CoV-2 revealed by SHAPE mutational profiling (SHAPEMaP) ^32^. In this approach, the authors performed *in vitro* refolding of RNA followed by probing with 2-methylnicotinic acid imidazolide. In our comparison, balanced lists of single and double stranded regions were used for the calculations: A confidence score of 10% indicates that we compared the SHAPE reactivity values of 3000 nucleotides associated with the highest *CROSS* scores (i.e., double stranded) and 3000 nucleotides associated with the lowest *CROSS* scores (i.e., single stranded). From low (10%) to high (0.1%) confidence scores, we observed that the predictive power, measured as the Area Under the Curve (AUC) of Receiver Operating Characteristics (ROC), increases monotonically reaching the value of 0.73 (the AUC is 0.74 for the 10 highest/lowest scores; **Fig. 1C**), which indicates that *CROSS* is able to reproduce SHAPEMaP in great detail.

We also assessed *CROSS* performances on structures of betacoronavirus 5’ and 3’ ends ^33–36^ (**Fig. 1D**). In this analysis, we used RFAM multiple sequence alignments of betacoronavirus 5’ and 3’ ends and relative consensus structures (RF03117 and RF03122) ^33–36^. We generated the 2D representation of nucleotide chains of consensus structures. We extracted the ‘secondary structure occupancy’, as defined in a previous work ^20^, and counted the contacts present around each nucleotide. Following the procedure used for the comparison with SHAPEMaP, different progressive cut-offs were used for ranking all the structures using balanced lists of single and double stranded regions: 10% indicates that we compared 600 nucleotides associated with the highest amount of contacts and 600 nucleotides associated with the lowest amount of contacts. From low (10%) to high (0.1%) confidence scores we observed that the AUC of ROC increases monotonically reaching the value of 0.75 (10 highest/lowest scores have an AUC of 0.78; **Fig. 1C**), which indicates that *CROSS* is able to identify known double and single stranded regions reported in great detail. We also tested the ability of *CROSS* to recognize specific secondary structures in representative cases for which we studied both the 3’ and 5’ ends: NC_006213 or Human coronavirus OC43 strain ATCC VR-759, NC_019843 or Middle East respiratory syndrome coronavirus, NC_026011 or Betacoronavirus HKU24 strain HKU24-R05005I, NC_001846 or Mouse hepatitis virus strain MHV-A59 C12 and NC_012936 or Rat coronavirus Parker (**Supp. Fig. 2**).

**Fig. 2.**
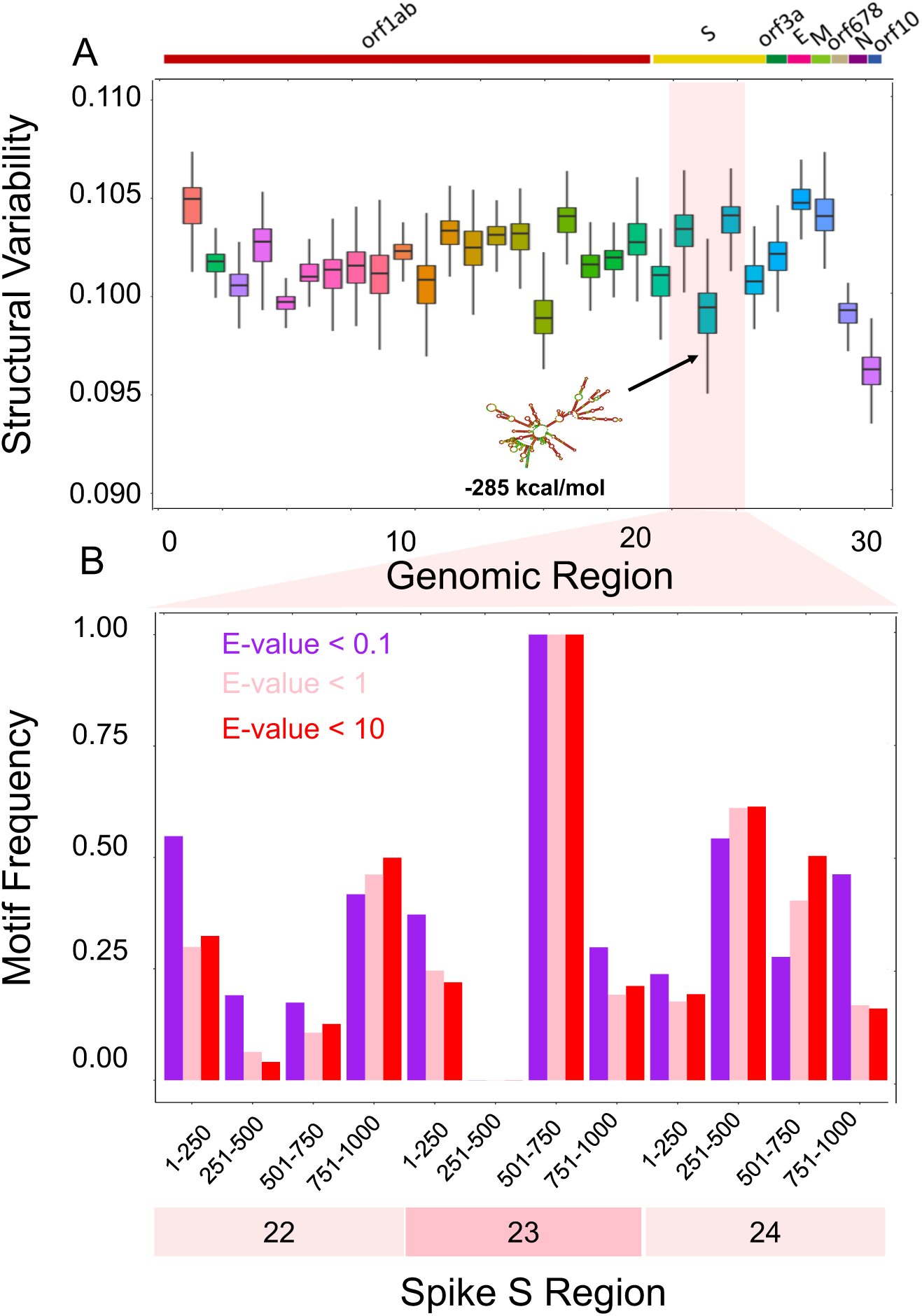
Structural comparisons of coronaviruses. **(A)** We employed the CROSSalign approach ^13,12^ to compare Wuhan strain MN908947.3 with other coronaviruses. One of the regions with the lowest structural variability encompasses nucleotides 22000-23000. The centroid structure and free energy computed with the Vienna method ^27^ are displayed. (**B**) We studied the the conservation of nucleotides 22000-23000 (fragment 23) and the adjacent regions using structural motives identified with RF-Fold algorithm of the RNAFramework suite ^30^ with CROSS as soft-constraint. We found that nucleotides 501-750 in fragment 23 are the ones with the highest number of matches for different found that nucleotides 501-750 within fragment 23 are the ones with the highest number of matches at confidence thresholds (E-values).

In summary, our analysis identifies several structural elements in SARS-CoV-2 genome ^11^. Different lines of experimental and computational evidence indicate that transcripts containing a large amount of double-stranded regions have a strong propensity to recruit proteins ^14^ and can act as scaffolds for protein assembly ^15,16^. We expected that the 5’ end attracts several host proteins because of the enrichment in secondary structure elements. The binding would not just involve proteins interacting with double-stranded regions. If a specific protein contact occurs in a loop at the end of a long RNA stem, the overall region is enriched in double-stranded nucleotides but the specific interaction takes place in a single-stranded element.

### Structural comparisons reveal that a spike S region of SARS-CoV-2 is conserved among coronaviruses

We employed *CROSS*align ^13^ to study the structural conservation of SARS-CoV-2 in different strains (**Materials and Methods**).

In our analysis, we compared the Wuhan strain MN908947.3 with 2040 coronaviruses (reduced to 267 sequences upon redundancy removal at 95% sequence similarity ^37^; **Fig. 2**; full data shown in **Supp. Fig. 1**). The 5’ end is highly variable. However, it is more structured in SARS-CoV-2 than other coronaviruses (average structural content of 0.56, indicating that 56% of the *CROSS* signal is >0). The 3’ end is less variable and slightly less structured (average structural content of 0.49). By contrast, the other coronaviruses have lower average structural content of 0.49 in the 5’ end and 0.42 in the 3’ end. One conserved region falls inside the Spike S genomic locus between nucleotides 22000 - 23000 and exhibits an intricate and stable secondary structure (RNA*fold* minimum free energy= -285kcal/mol)^27^. High conservation of a structured region suggests a functional activity that is relevant for host infection.

To demonstrate the conservation of nucleotides 22000-23000 (fragment 23), we divided this region and the adjacent ones (nucleotides 21000-22000 and 23000-24000) into sub-fragments. We then used the *RF-Fold* algorithm of the *RNAFramework* suite ^30^ to fold the different sub-regions using *CROSS* predictions as soft-constraints. The structural motives identified with this procedure were employed to build covariance models (CMs) that were then searched in our set of coronaviruses using the ‘Infernal’ package ^38^. We found that nucleotides 501-750 within fragment 23 have the highest number of matches for different confidence thresholds, implying a higher chance of sequence and structure conservation across coronaviruses (E-values of 10,1, 0.1; **Fig. 2B**). We specifically counted the matches falling in the Spike S region (+/-1000 nucleotides to take into account the division of the genome into fragments; **Supp. Table 1)**. For the large majority of annotated sequences, we found a match falling in the Spike S region (239 genomes out of 246, of which 161 with E-value below 0.1) This further emphasizes the conservation of the region in exam.

### Sequence and structural comparisons among SARS-CoV-2 strains

To better investigate the sequence conservation of SARS-CoV-2, we compared >60 strains isolated from different countries during the pandemic (including China, USA, Japan, Taiwan, India, Brazil, Sweden, and Australia; data from NCBI and in VIPR www.viprbrc.org; **Materials and Methods**). Our analysis aims to determine the relationship between structural content and sequence conservation.

Using *Clustal W* for multiple sequence alignments ^39^, we observed general conservation of the coding regions (**Fig. 3A**). The 5’ and 3’ ends show high variability due to experimental procedures of the sequencing and are discarded in this analysis ^40^. One highly conserved region is between nucleotides 22000 - 23000 in the Spike S genomic locus, while sequences up-and downstream are variable (red bars in **Fig. 3A**). We then used *CROSSalign* ^13^ to compare the structural content (**Materials and Methods**). High variability of structure is observed for both the 5’ and 3’ ends and for nucleotides 21000 - 22000 as well as 24000 - 25000, associated with the Spike S region (purple bars in **Fig. 3A**). The rest of the regions are significantly conserved at a structural level (p-value < 0.0001; Fisher’s test).

**Fig. 3.**
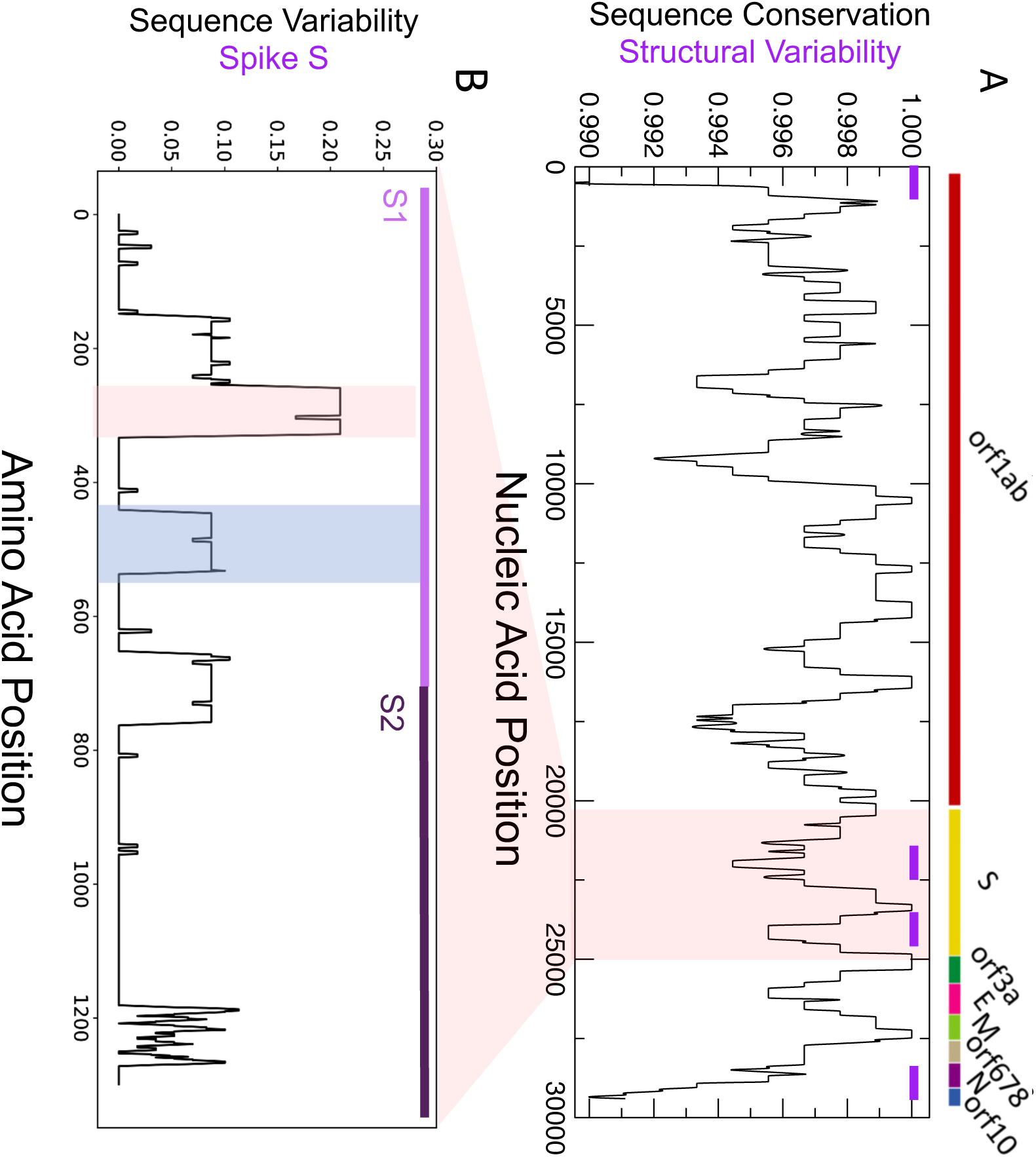
Sequence and structural comparison of human SARS-CoV-2 strains. (**A**) Strong sequence conservation (Clustal W multiple sequence alignments ^49^) is observed in coding regions, including the region between nucleotides 22000 and 23000 of spike S protein. High structural variability (red bars on top) is observed for both the UTRs and for nucleotides between 21000 and 22000 as well as 24000 and 25000, associated with the S region. The rest of the regions are significantly conserved at a structural level. (**B**) The sequence variability (Shannon entropy computed on T-Coffee multiple sequence alignments ^43^) in the spike S protein indicate conservation between amino-acids 460 and 520 (blue box) binding to the host receptor angiotensin-converting enzyme 2 ACE2. The region encompassing amino-acids 243 and 302 is highly variable and is implicated in sialic acids in MERS-CoV (red box). The S1 and S2 domains of Spike S protein are displayed.

We note that sequence conservation (**Fig. 3A**) and secondary structure profiles (**Fig. 1A**) are statistically related. Following the analysis to compare *CROSS* and SHAPE scores, we selected balanced groups of nucleotides with the highest and lowest sequence conservation and measured their single and double stranded content: a conservation score of 1% indicates that we compared 300 nucleotides with the highest sequence similarity and 300 nucleotides with the lowest sequence similarity. At conservation score of 1% (or less stringent threshold of 10%), the match between similarity and structure, measured as the AUC of ROC is 0.76 (or 0.60, respectively). The association is statistically significant: shuffling the sequence conservation profiles, the empirical p-values are < 0.02 (at both 10% and 1% conservation scores).

We also compared protein sequences coded by the Spike S genomic locus (NCBI reference QHD43416) and found that both sequence (**Fig. 3A**) and structure (**Fig. 2**) of nucleotides 22000 - 23000 are highly conserved. The region corresponds to amino acids 330-500 that contact the host receptor angiotensin-converting enzyme 2 (ACE2) ^41^ promoting infection and provoking lung injury ^24,42^. By contrast, the region upstream of the binding site receptor ACE2 and located in correspondence to the minimum of the structural profile at around nucleotides 22500-23000 (**Fig. 1**) is highly variable ^43^, as indicated by *T-coffee* multiple sequence alignments ^43^ (**Fig. 3A**). This part of the Spike S region corresponds to amino acids 243-302 that in MERS-CoV binds to sialic acids regulating infection through cell-cell membrane fusion (**Fig. 3B;** see related manuscript by E. Milanetti *et al*.) ^10–44,45^.

Our analysis suggests that the structural region between nucleotides 22000 and 23000 of Spike S region is conserved among coronaviruses (**Fig. 2**) and that the binding site for ACE2 has poor variation in human SARS-CoV-2 strains (**Fig. 3B**). By contrast, the region upstream, which has propensity to bind sialic acids ^10–44,45^, showed poor structural content and high variability (**Fig. 3B**).

### Analysis of human interactions with SARS-CoV-2 identifies proteins involved in viral replication

In order to obtain insights on how the virus replicates in human cells, we predicted SARS-CoV-2 interactions with the whole RNA-binding human proteome. Following a protocol to study structural conservation in viruses ^13^, we first divided the Wuhan sequence in 30 fragments of 1000 nucleotides each moving from the 5’ to 3’ end and then calculated the protein-RNA interactions of each fragment with *cat*RAPID *omics* (3340 canonical and putative RNA-binding proteins, or RBPs, for a total 102000 interactions) ^18^. Proteins such as Polypyrimidine tract-binding protein 1 PTBP1 (Uniprot P26599) showed the highest interaction propensity (or Z-score; **Materials and Methods**) at the 5’ end while others such as Heterogeneous nuclear ribonucleoprotein Q HNRNPQ (O60506) showed the highest interaction propensity at the 3’end, in agreement with previous studies on coronaviruses (**Fig. 4A**) ^46^.

**Fig. 4.**
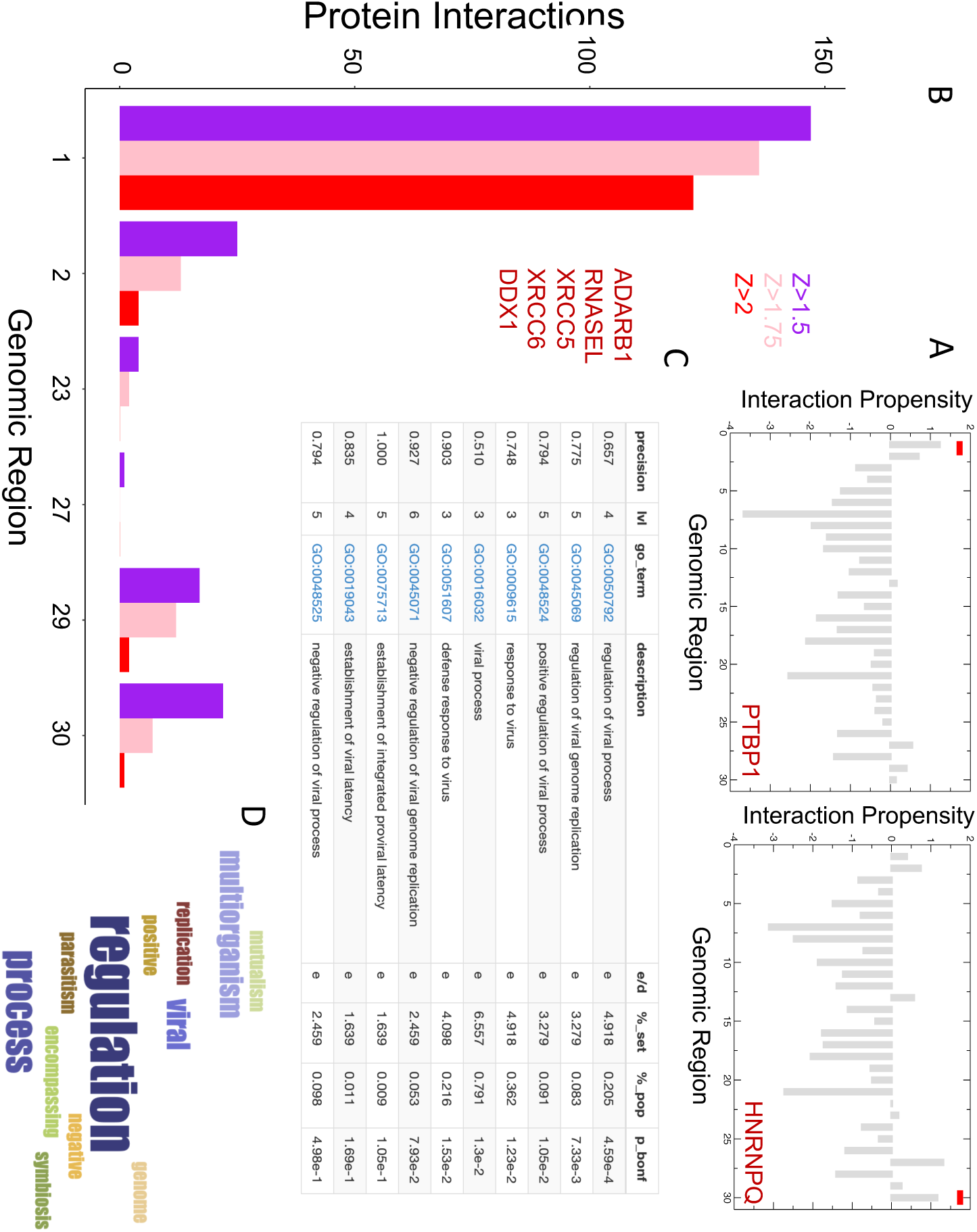
Predictions of protein interactions with SARS-CoV-2 RNA. (**A**) In agreement with studies on coronaviruses ^46^, PTBP1 shows the highest interaction propensity at the 5’ and HNRNPQ at 3’ (indicated by red bars). (**B**) Number of RBP interactions for different SARS-CoV-2 regions (colours indicate catRAPID ^18,19^ confidence levels: Z=1.5 or low Z=1.75 or medium and Z=2.0 or high; regions with scores lower than Z=1.5 are omitted); (**C**) Enrichment of viral processes in the 5’ of SARS-CoV-2 (precision = term precision calculated from the GO graph structure lvl = depth of the term; go_term = GO term identifier, with link to term description at AmiGO website; description = label for the term; e/d = enrichment / depletion compared to the population; %_set = coverage on the provided set; %_pop = coverage of the same term on the population; p_bonf = p-value of the enrichment. To correct for multiple testing bias, use Bonferroni correction) ^49^; (**D**) Viral processes are the third largest cluster identified in our analysis;

For each fragment, we predicted the most significant interactions by filtering according to the Z score. We used three different thresholds in ascending order of stringency: Z ≥ 1.50, 1.75 and 2 respectively and we removed from the list the proteins that were predicted to interact promiscuously with more than one fragment. Fragment 1 corresponds to the 5’ end and is the most contacted by RBPs (around 120 with Z≥2 high-confidence interactions; **Fig. 4B**), which is in agreement with the observation that highly structured regions attract a large number of proteins ^14^. Indeed, the 5’ end contains multiple stem loop structures that control RNA replication and transcription ^47,48^. By contrast, the 3’ end and fragment 23 (Spike S), which are still structured but to a lesser extent, attract fewer proteins (10 and 5, respectively) and fragment 20 (between Orf1ab and Spike S) that is predicted to be unstructured, does not have predicted binding partners.

The interactome of each fragment was analysed using *clever*GO, a tool for Gene Ontology (GO) enrichment analysis ^49^. Proteins interacting with fragments 1, 2 and 29 were associated with annotations related to viral processes (**Fig. 4C; Supp. Table 2**). Considering the three thresholds applied (**Materials and Methods**), we found 23 viral proteins (including 2 pseudogenes), for fragment 1, 2 proteins for fragment 2 and 11 proteins for fragment 29 (**Fig. 4D**). Among the high-confidence interactors of fragment 1, we discovered RBPs involved in positive regulation of viral processes and viral genome replication, such as double-stranded RNA-specific editase 1 ADARB1 (Uniprot P78563), 2-5A-dependent ribonuclease RNASEL (Q05823) and 2-5-oligoadenylate synthase 2 OAS2 (P29728; **Fig. 5A**). Interestingly, 2-5-oligoadenylate synthase 2 OAS2 has been reported to be upregulated in human alveolar adenocarcinoma (A549) cells infected with SARS-CoV-2 (log fold change of 4.2; p-value of 10^−9^ and q-value of 10^−6^) ^50^. While double-stranded RNA-specific adenosine deaminase ADAR (P55265) is absent in our library due to its length that does not meet *cat*RAPID *omics* requirements ^18^, the *omiXcore* extension of the algorithm specifically developed for large molecules ^51^ attributes the same binding propensity to both ADARB1 and ADAR, thus indicating that the interactions with SARS-CoV-2 are likely to occur (**Materials and Methods**). Moreover, experimental works indicate that the family of ADAR deaminases is active in bronchoalveolar lavage fluids derived from SARS-CoV-2 patients ^52^ and is upregulated in A549 cells infected with SARS-CoV-2 (log fold change of 0.58; p-value of 10^−8^ and q-value of 10^−5^) ^50^.

**Fig. 5.**
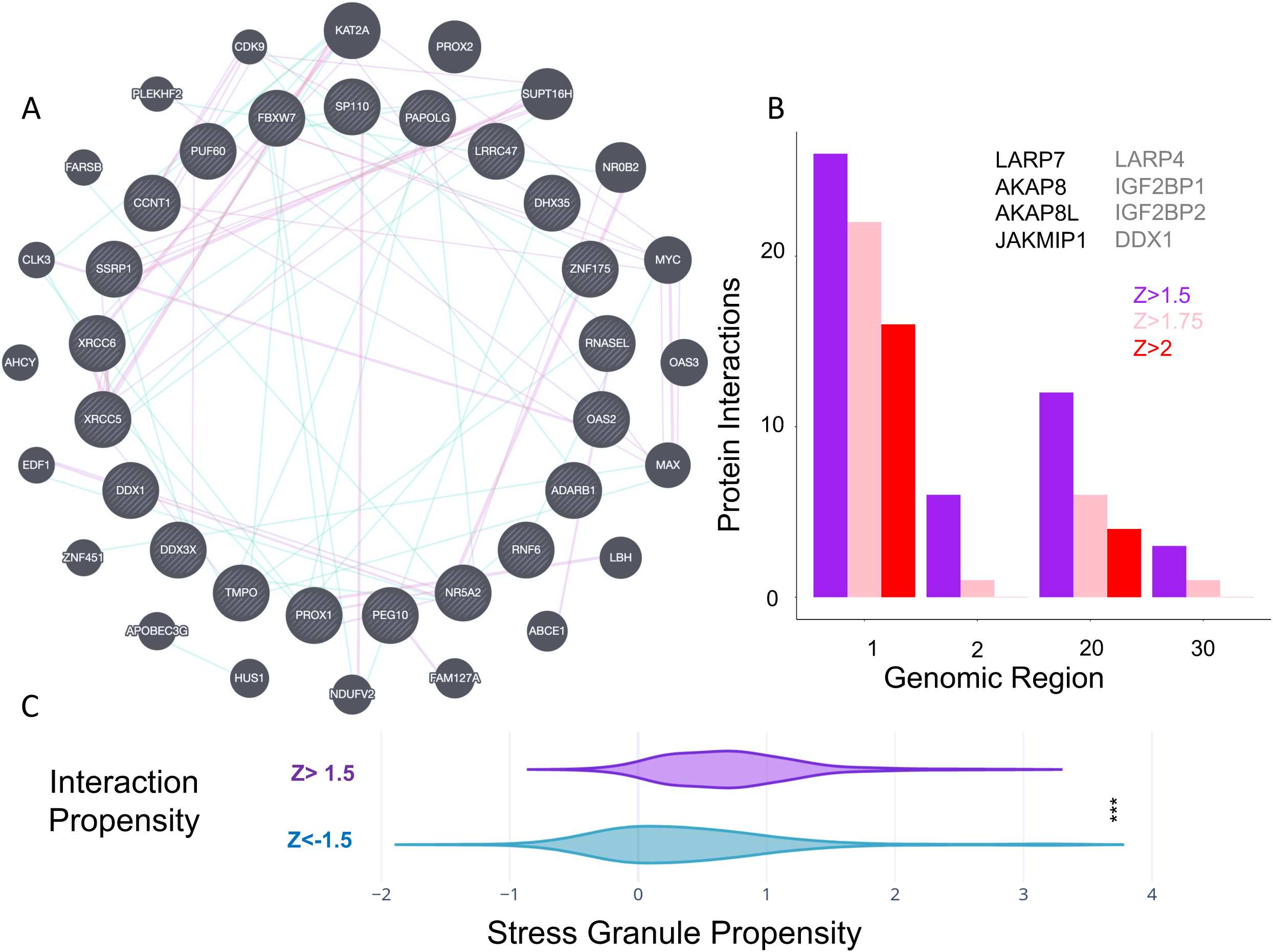
Characterization of protein interactions with SARS-CoV-2 RNA. (**A**) Protein interaction network of SARS-CoV-2 5’ end (inner circle) and associations with other human genes retrieved from literature (blue: genetic associations; purple: physical associations); (**B**) Number of RBP interactions identified by Gordon et al. ^67^ and Schmidt et al. ^73^ for different SARS-CoV-2 regions. Representative cases are shown in black (Gordon et al. ^67^) and grey (Schmidt et al. ^73^). (**C**) Proteins binding to the 5’ with Z score ≥ 1.5 show high propensity to accumulate in stress-granules (same number of proteins with Z score < -1.5 are used in the comparison; *** p-value<0.0001; Kolmogorov-Smirnoff).

We also identified proteins related to the establishment of integrated proviral latency, including X-ray repair cross-complementing protein 5 XRCC5 (P13010) and X-ray repair cross-complementing protein 6 XRCC6 (P12956; **Fig. 5A**). In accordance with our calculations, comparison of A549 cells responses to SARS-CoV-2 and respiratory syncytial virus, indicates upregulation of XRRC6 in SARS-CoV-2 (log fold-change of 0.92; p-value of 0.006 and q-value of 0.23) ^50^. Moreover, previous evidence suggests that the binding of XRCC6 takes places at the 5’ end of SARS-CoV-2, thus giving further support to our predictions ^53^. Nucleolin NCL (P19338), a protein known to be involved in coronavirus processing, was also predicted to bind tightly to the 5’ end (**Supp. Table 2**) ^54^.

Importantly, we found proteins related to defence response to viruses, such as ATP-dependent RNA helicase DDX1 (Q92499), that are involved in negative regulation of viral genome replication. Some DNA-binding proteins such as Cyclin-T1 CCNT1 (O60563), Zinc finger protein 175 ZNF175 (Q9Y473) and Prospero homeobox protein 1 PROX1 (Q92786) were included because they could have potential RNA-binding ability (**Fig. 5A**) ^55^. As for fragment 2, we found two canonical RBPs: E3 ubiquitin-protein ligase TRIM32 (Q13049) and E3 ubiquitin-protein ligase TRIM21 (P19474), which are listed as negative regulators of viral release from host cell, negative regulators of viral transcription and positive regulators of viral entry into host cells. Among these genes, DDX1 (log fold change of 0.36; p-value of 0.007 and q-value of 0.23) and TRIM21 (log fold change of 0.44; p-value of 0.003 and q-value of 0.18) are also upregulated in A549 cells infected with SARS-CoV-2 ^50^. Ten of the 11 viral proteins detected for fragment 29 are members of the Gag polyprotein family, that perform different tasks during HIV assembly, budding, and maturation. More than just scaffold elements, these proteins are elements that accompany viral and host proteins as they traffic to the cell membrane (**Supp. Table 2**) ^56^. Finally, among the RBPs with the highest interaction propensity for fragment 23, we found nucleosome assembly protein 1-like 1 NAP1L1 and E3 ubiquitin-protein ligase makorin-1 MKRN1, which could have an effect on the regulation of cell proliferation.

Analysis of functional annotations carried out with *GeneMania* ^57^ revealed that proteins interacting with the 5’ of SARS-CoV-2 RNA are associated with regulatory pathways involving NOTCH2, MYC and MAX that have been previously connected to viral infection processes (**Fig. 5A**) ^58,59^. Interestingly, some proteins, including DDX1, CCNT1 and ZNF175 for fragment 1 and TRIM32 for fragment 2, have been shown to be necessary for HIV functions and replication inside the cell, as well as SARS-CoV-1. DDX1 has been shown to enable the switch from discontinuous to continuous transcription in SARS-CoV-1 infection and its knockdown reduced the number of longer sub-genomic mRNA (sgmRNA) through interaction with the SARS-CoV-1 nucleocapsid protein N ^60^ and Nsp14 ^61^. It functions as a bidirectional helicase, which distinguishes it from the coronaviruses helicases, which can only unwind RNA in the 5’ to 3’ direction ^62^, a very important function in highly structured RNA such SARS-CoV-2. DDX1 is also required for HIV-1 Rev as well as for avian coronavirus IBV replication and it binds to the RRE sequence of HIV-1 RNAs ^1–3^, while CCNT1 binds to 7SK snRNA and regulates transactivation domain of the viral nuclear transcriptional activator, Tat ^65,66^.

### Analyses of SARS-CoV-2 proteins interactomes reveal common protein targets

Recently, Gordon *et al*. reported a list of human proteins binding to Open Reading Frames (ORFs) translated from SARS-CoV-2 ^67^. Identified through affinity purification followed by mass spectrometry quantification, 332 proteins from HEK-293T cells interact with viral ORF peptides. By selecting 274 proteins binding at the 5’ with Z score ≥1.5 (**Supp. Table 2**), of which 140 are exclusively interacting with fragment 1 (**Fig. 4B**), we found that 8 are also reported in the list by Gordon *et al*. ^67^, which indicates significant enrichment (representation factor of 2.5; p-value of 0.02; hypergeometric test with human proteome in background). The fact that our list of protein-RNA binding partners contains elements identified also in the protein-protein network analysis is not surprising, as ribonucleoprotein complexes evolve together ^14^ and their components sustain each other through different types of interactions ^16^.

We note that out of 332 interactions, 60 are RBPs (as reported in Uniprot), which represents a considerable fraction (i.e., 20%), considering that there are around 1500 RBPs in the human proteome (i.e., 6%). Comparing the RBPs present in Gordon *et al*. ^67^ and those present in our list (79 RBP annotated in Uniprot), we found an overlap of 6 proteins (representation factor = 26.5; p-value < 10^−8^; hypergeometric test), including: Janus kinase and microtubule-interacting protein 1 JAKMIP1 (Q96N16), A-kinase anchor protein 8 AKAP8 (O43823) and A-kinase anchor protein 8-like AKAP8L (Q9ULX6), which in case of HIV-1 infection is involved as a DEAD/H-box RNA helicase binding ^69^, signal recognition particle subunit SRP72 (O76094), binding to the 7S RNA in presence of SRP68, La-related protein 7, LARP7 (Q4G0J3) and La-related protein 4B LARP4B (Q92615), which are part of a system for transcriptional regulation acting by means of the 7SK RNP system (**Fig. 5B; Supp. Table 3**). We speculate that sequestration of these elements is orchestrated by a viral program aiming to recruit host genes ^71^. LARP7 is also upregulated in A549 cells infected with SARS-CoV-2 (log fold change of 0.48; p-value of 0.006 and q-value of 0.23) ^50^.

Moreover, by directly analysing the RNA interaction potential of all the 332 proteins by Gordon *et al*. ^67^, *cat*RAPID identified 38 putative binders at the 5’ end (Z score ≥ 1.5; 27 occurring exclusively in the 5’ end and not in other regions of the RNA) ^18^, including Serine/threonine-protein kinase TBK1 (Q9UHD2), among which 10 RBPs (as reported in Uniprot) such as: Splicing elements U3 small nucleolar ribonucleoprotein protein MPP10 (O00566) and Pre-mRNA-splicing factor SLU7 (O95391), snRNA methylphosphate capping enzyme MEPCE involved in negative regulation of transcription by RNA polymerase II 7SK (Q7L2J0) ^72^, Nucleolar protein 10 NOL10 (Q9BSC4) and protein kinase A Radixin RDX (P35241; in addition to those mentioned above; **Supp. Table 3**).

Using the liver cell line HuH7 a recent experimental study by Schmidt *et al*. ^73^. identified SARS-CoV-2 RNA associations within the human host ^73^. Through the RAP-MS approach, 571 interactions were detected, of which 250 are RBPs (as reported in Uniprot) ^73^.

In common with our library we found an overlap of 148 proteins. We compared predicted (as released in March 2020) and experimentally-validated interactions employing balanced lists of high-affinity (high fold-change with respect to RNA Mitochondrial RNA Processing Endoribonuclease RMRP) and low-affinity (low fold-change with respect to RNA Mitochondrial RNA Processing Endoribonuclease RMRP) associations: a confidence score of 25% indicates that we compared the interaction scores of 35 proteins with the highest fold-change values and 35 interactions associated with the lowest fold-change values. From low (25%) to high (5%) confidence scores, we observed that the predictive power, measured as the Area Under the Curve (AUC) of Receiver Operating Characteristics (ROC), increases monotonically reaching the value of 0.99 (the AUC is 0.72 for 25% confidence score; **Supp. Fig. 3**), which indicates strong agreement between predictions and experiments. In addition to DDX1 and DDX3X (O00571), other interactions corresponding to *cat*RAPID scores > 1.5 and fold-change > 1 include Insulin-like growth factor 2 mRNA-binding protein 1 IGF2BP1 (Q9Y6M1), Insulin-like growth factor 2 mRNA-binding protein 2 IGF2BP2 2 (Q9Y6M1) and La-related protein 4 LARP4 (Q71RC2; also in Gordon et al. ^67^).

By directly analysing RNA interactions of all the 571 proteins by Schmidt *et al*. ^73^, *cat*RAPID identified 18 strong RBP binders at the 5’ end (Z score ≥ 1.5; fold-change>1; p-value of 0.008 computed with respect to all the interactions; Fisher exact test; **Supp. Table. 4**), including Helicase MOV-10 (Q9HCE1), Cold shock domain-containing protein E1 CSDE1 (O75534), Staphylococcal nuclease domain-containing protein 1 SND1 (Q7KZF4), Pumilio homolog 1 PUM1 (Q14671), and La-related protein 1 LARP1 (Q6PKG0), among other interactions (**Supp. Table 4**).

### The 5’ end is enriched in host interactions implicated in other viral infections

In the list of 274 proteins binding to the 5’ end (fragment 1) with Z score ≥1.5, we found 10 hits associated with HIV (**Supp. Table 5**), which represents a significant enrichment (p-value=0.0004; Fisher’s exact test), considering that the total number of HIV-related proteins is 35 in the whole *cat*RAPID library (3340 elements). The complete list of proteins includes ATP-dependent RNA helicase DDX1 (Q92499), ATP-dependent RNA helicase DDX3X (O00571 also involved in Dengue and Zika Viruses), Tyrosine-protein kinase HCK (P08631, nucleotide binding), Arf-GAP domain and FG repeat-containing protein 1 (P52594), Double-stranded RNA-specific editase 1 ADARB1 (P78563), Insulin-like growth factor 2 mRNA-binding protein 1 IGF2BP1 (Q9NZI8), A-kinase anchor protein 8-like AKAP8L (Q9ULX6; its partner AKAP8 is also found in Gordon *et al*. ^67^) Cyclin-T1 CCNT1 (O60563; DNA-binding) and Forkhead box protein K2 FOXK2 (Q01167; DNA-binding; **Fig. 4B** and **Fig. 5A; Supp. Table 5**).

Smaller enrichments were found for proteins related to Hepatitis B virus (HBV; p-value=0.01; 3 hits out of 7 in the whole *cat*RAPID library; Fisher’s exact test), including Nuclear receptor subfamily 5 group A member 2 NR5A2 (DNA-binding; O00482), Interferon-induced, double-stranded RNA-activated protein kinase EIF2AK2 (P19525), and SRSF protein kinase 1 SRPK1 (Q96SB4) as well as Influenza A (p-value=0.03; 2 hits out of 4; Fisher’s exact test), including Synaptic functional regulator FMR1 (Q06787) and RNA polymerase-associated protein RTF1 homologue (Q92541; **Supp. Table 5**). By contrast, no significant enrichments were found for other viruses such as for instance Ebola.

Very importantly, specific chemical compounds have been developed to interact with HIV-and HVB-related proteins. The list of HIV-related targets reported in ChEMBL ^74^ includes ATP-dependent RNA helicase DDX1 (CHEMBL2011807), ATP-dependent RNA helicase DDX3X (CHEMBL2011808), Cyclin-T1 CCNT1 (CHEMBL2348842), and Tyrosine-protein kinase HCK (CHEMBL2408778), among other targets. In addition, HVB-related targets are Nuclear receptor subfamily 5 group A member 2 NR5A2 (CHEMBL3544), Interferon-induced, double-stranded RNA-activated protein kinase EIF2AK2 (CHEMBL5785) and SRSF protein kinase 1 SRPK1 (CHEMBL4375). We hope that this list can be the starting point for further pharmaceutical studies.

### Phase-separating proteins are enriched in the 5’ end interactions

As SARS-CoV-2 represses host gene expression through a number of unknown mechanisms, sequestration of cell transcription machinery elements could be exploited to alter biological pathways in the host cell. A number of proteins identified in our *cat*RAPID calculations have been previously reported to coalesce in large ribonucleoprotein assemblies similar to stress granules. Among these proteins, we found double-stranded RNA-activated protein kinase EIF2AK2 (P19525), Nucleolin NCL (P19338), ATP-dependent RNA helicase DDX1 (Q92499), Cyclin-T1 CCNT1 (O60563), signal recognition particle subunit SRP72 (O76094), LARP7 (Q4G0J3) and La-related protein 4B LARP4B (Q92615) as well as Polypyrimidine tract-binding protein 1 PTBP1 (P26599) and Heterogeneous nuclear ribonucleoprotein Q HNRNPQ (O60506) ^75^. To further investigate the propensity of these proteins to phase separate, we used the *cat*GRANULE algorithm (**Materials and Methods**) ^76^. We found that the 274 proteins binding to the 5’ end (fragment 1) with Z score ≥1.5 are highly prone to accumulate in assemblies similar to stress-granules (274 proteins with the lowest Z score are used in the comparison; p-value<0.0001; Kolmogorov-Smirnoff; **Fig. 5C; Supp. Table 6**).

Supporting this hypothesis, DDX1 and CCNT1 have been shown to condense in membrane-less organelles such as stress granules ^77–79^ that are the direct target of RNA viruses ^80^. DDX1 is also the primary component of distinct nuclear *foci* ^81^, together with factors associated with pre-mRNA processing and polyadenylation. Similarly, SRP72, LARP7 and LARP4B proteins have been found to assemble in stress granules ^82–83,75^. A recent work also suggests that the binding of LARP4 and XRCC6 takes places at the 5’ end of SARS-CoV-2 and contributes to SARS-CoV-2 phase separation ^53^. Moreover, emerging evidence indicates that the SARS-CoV-2 nucleocapsid protein N has a strong phase separation propensity that is modulated by the viral genome ^53–84,85^ and can enter into host cell protein condensates ^84^, suggesting a possible mechanism of cell protein sequestration. Notably, *ca*tGRANULE does predict that nucleocapsid protein N is the viral protein with highest propensity to phase separate ^86^.

Our observations are particularly relevant because RNA viruses are known to antagonize stress granules formation ^80^. Indeed, the role of stress granules and processing bodies in translation suppression and RNA decay have impact on virus replication ^87^.

## CONCLUSIONS

Our study is motivated by the need to identify molecular mechanisms involved in Covid-19 spreading. Using advanced computational approaches, we investigated the structural content of SARS-CoV-2 genome and predicted human proteins that bind to it.

We employed *CROSS* ^13,12^ to compare the structural properties of around 2000 coronaviruses and identified elements conserved in SARS-CoV-2 strains. The regions containing the highest amount of structure are the 5’ end as well as glycoproteins spike S and membrane M.

We found that the Spike S protein domain encompassing amino acids 460-520 is conserved across SARS-CoV-2 strains. This result suggests that Spike S must have evolved to specifically interact with its host partner ACE2 ^41^ and mutations increasing the binding affinity should be highly infrequent. As nucleic acids encoding for this region are enriched in double-stranded content, we speculate that the structure might attract host regulatory elements, such as nucleosome assembly protein 1-like 1 NAP1L1 and E3 ubiquitin-protein ligase makorin-1 MKRN1, further constraining its variability. The fact that this region of the Spike S region is highly conserved among all the analysed SARS-CoV-2 strains suggests that a specific drug could be designed to prevent interactions within the host.

The highly variable region at amino acids 243-302 in spike S protein corresponds to the binding site of sialic acids in MERS-CoV ^7–10,45^ and could play a role in infection ^44^. The fact that the binding region is highly variable suggests different affinities for sialic acid-containing oligosaccharides and polysaccharides such as heparan sulfate, which provides clues on the specific responses in the human population. At present, a glycan microarray technology indicated that SARS-CoV-2 Spike S binds more tightly to heparan sulfate than sialic acids ^88^.

Using *cat*RAPID ^18,19^ we computed >100000 protein interactions with SARS-CoV-2 and found previously reported interactions such as Polypyrimidine tract-binding protein 1 PTBP1, Heterogeneous nuclear ribonucleoprotein Q HNRNPQ and Nucleolin NCL ^54^. In addition, we discovered that the highly structured region at the 5’end has the largest number of protein partners including ATP-dependent RNA helicase DDX1, which was previously reported to be essential for HIV-1 and coronavirus IBV replication ^63,64^, and the double-stranded RNA-specific editases ADAR and ADARB1, which catalyse the hydrolytic deamination of adenosine to inosine. Other predicted interactions are XRCC5 and XRCC6 members of the HDP-RNP complex associating with ATP-dependent RNA helicase DHX9 ^90^ as well as and 2-5A-dependent ribonuclease RNASEL and 2-5-oligoadenylate synthase 2 OAS2 that control viral RNA degradation ^91,92^. Interestingly, DDX1, XRCC6 and OAS2 were found upregulated in human alveolar adenocarcinoma cells infected with SARS-CoV-2 ^50^ and DDX1 knockdown has been shown to reduce the number of sgmRNA in SARS-CoV-1 infected cells ^60^. In agreement with our predictions, recent experimental work indicates that the family of ADAR deaminases is active in bronchoalveolar lavage fluids derived from SARS-CoV-2 patients ^52^.

Comparison with protein-RNA interactions detected in the liver cell line HuH7 ^73^ shows agreement with our predictions. We note that the experiments have been carried out 24 hours after infection ^73^, which indicates that the protein interaction landscape might have changed with respect to the early events of replication. Yet, the accordance with our calculations indicates participation of elements involved in controlling RNA processing and editing (DDX1, DDX3X) and translation (IGF2BP1 and IGF2BP2), although proteins such as ADAR and XRCC5 were reported to have poorer binding capacity ^73^.

A significant overlap exists with the list of protein interactions reported by Gordon *et al*. ^67^, and among the candidate partners we identified AKAP8L, involved as a DEAD/H-box RNA helicase binding protein involved in HIV infection ^69^. In general, proteins associated with retroviral replication are expected to play different roles in SARS-CoV-2. As SARS-CoV-2 massively represses host gene expression ^71^, we hypothesize that the virus hijacks host pathways by recruiting transcriptional and post-transcriptional elements interacting with polymerase II genes and splicing factors such as for instance A-kinase anchor protein 8-like AKAP8L and La-related protein 7 LARP7. In concordance with our predictions LARP7 has been reported to be upregulated in human alveolar adenocarcinoma cells infected with SARS-CoV-2 ^50^. The link to proteins previously studied in the context of HIV and other viruses, if further confirmed, is particularly relevant for the repurposing of existing drugs ^74^.

The idea that SARS-CoV-2 sequesters different elements of the transcriptional machinery is particularly intriguing and is supported by the fact that a large number of proteins identified in our screening are found in stress granules ^75^. Indeed, stress granules protect the host innate immunity and are hijacked by viruses to favour their own replication ^87^. Moreover, as coronaviruses transcription uses discontinuous RNA synthesis that involves high-frequency recombination ^54^, it is possible that pieces of the viruses resulting from a mechanism called defective interfering RNAs ^93^ could act as scaffold to attract host proteins ^14,15^. In agreement with our hypothesis, it has been very recently shown that he coronavirus nucleocapsid protein N can form protein condensates based on viral RNA scaffold and can merge with the human cell protein condensates ^84^, which provides a potential mechanism of host protein sequestration.

## Acknowledgements

The authors would like to thank Dr. Mattia Miotto, Dr Lorenzo Di Rienzo, Dr. Alexandros Armaos, Dr. Alessandro Dasti and Dr. Claudia Giambartolomei for discussions. We are particularly grateful to Prof. Annalisa Pastore for critical reading, Dr. Gilles Mirambeau for the RT vs RdRP analysis, Dr. Andrea Cerase for the discussing on stress granules and Dr. Roberto Giambruno for pointing to PTBP1 and HNRNPQ experiments.

The research leading to these results has been supported by European Research Council (RIBOMYLOME_309545 and ASTRA_855923), the H2020 projects IASIS_727658 and INFORE_825080, the Spanish Ministry of Economy and Competitiveness BFU2017-86970-P as well as the collaboration with Peter St. George-Hyslop financed by the Wellcome Trust.

## Contributions

GGT and RDP conceived the study. AV carried out *cat*RAPID analysis of protein interactions, RDP calculated *CROSS* structures of coronaviruses, GGT, MM and EM performed and analysed sequence alignments, JR, EZ and EB analysed the prediction results. AV, RDP and GGT wrote the paper.

**The authors do not have conflicts of interests**.

## MATERIALS AND METHODS

### Structure prediction

We predicted the secondary structure of transcripts using *CROSS* (Computational Recognition of Secondary Structure) ^13,12^. The algorithm predicts the structural profile (single-and double-stranded state) at single-nucleotide resolution using sequence information only and without sequence length restrictions (scores > 0 indicate double stranded regions). We used the *Vienna* RNA Package ^27^ to further investigate the RNA secondary structure of minima and maxima identified with *CROSS* ^13^.

*CROSS alive* was employed to predict SARS-CoV-2 secondary structure *in vivo* ^28^. *CROSS alive* (m6A+ fast option) predicts long range interactions and can identify pseudoknots of 50-100 nucleotides. The *RF-Fold* algorithm of the *RNAFramework* suite ^28^ was used to identify pseudoknots in SARS-CoV-2. In this analysis, the partition function was calculated using CROSS calculations as soft-constraints. RNA was then folded employing *Vienna* RNA Package ^27^ and pseudo-knotted bases were hard-constrained to be single-stranded.

MN908947 predictions are available at http://crg-webservice.s3.amazonaws.com/submissions/2020-05/270257/output/index.html?unlock=fd65439e7b (*CROSS*) and also http://crg-webservice.s3.amazonaws.com/submissions/2020-05/271372/output/index.html?unlock=1de1d3a54a (*CROSS alive*).

### Structural conservation

We used *CROSSalign* ^13,12^, an algorithm based on Dynamic Time Warping (DTW), to check and evaluate the structural conservation between different viral genomes ^13^. *CROSSalign* was previously employed to study the structural conservation of ∼5000 HIV genomes. SARS-CoV-2 fragments (1000 nt, not overlapping) were searched inside other complete genomes using the OBE (open begin and end) module, in order to search a small profile inside a larger one. The lower the structural distance, the higher the structural similarities (with a minimum of 0 for almost identical secondary structure profiles). The significance is assessed as in the original publication ^13^.

The Infernal package (version 1.1.3) was employed to build covariance models (CMs) for fragments 22,23 and 24 ^38^. The package was then used to search for sequence and structural similarities among RNAs in our database (267 representative sequences), which allows to identify a series of matches below a specific E-value threshold (0.1, 1 and 10). The analysis shows agreement with *CROSSalign* ^13,12^ results. The minimum and maximum number of identified motifs were 224 and 4878 (E-value of 10), 136 and 3093 (E-value of 1) and 94 and 1060 (E-value of 0.1). The motifs in Spike S region were counted for annotated coronaviruses (239 genomes out of 246, of which 161 within E-value of 0.1).

### Sequence collection

The FASTA sequences of the complete genomes of SARS-CoV-2 were downloaded in April 2020 from Virus Pathogen Resource (VIPR; www.viprbrc.org), for a total of 62 strains. Regarding the other coronaviruses, the sequences were downloaded in April 2020 from NCBI selecting only complete genomes, for a total of 2040 genomes. The reference Wuhan sequence with available annotation (EPI_ISL_402119) was downloaded from Global Initiative on Sharing All Influenza Data in April 2020. (GISAID https://www.gisaid.org/).

### Protein-RNA interaction prediction

Interactions between each fragment of target sequence and the human proteome were predicted using *cat*RAPID *omics* ^18,19^, an algorithm that estimates the binding propensity of protein-RNA pairs by combining secondary structure, hydrogen bonding and van der Waals contributions. As reported in a recent analysis of about half a million of experimentally validated interactions ^31^, the algorithm is able to separate interacting vs non-interacting pairs with an area under the ROC curve of 0.78. The complete list of interactions between the 30 fragments and the human proteome is available at http://crg-webservice.s3.amazonaws.com/submissions/2020-03/252523/output/index.html?unlock=f6ca306af0. The output then is filtered according to the Z-score column, which is the interaction propensity normalised by the mean and standard deviation calculated over the reference RBP set (http://s.tartaglialab.com/static_files/shared/faqs.html#4). We used three different thresholds in ascending order of stringency: Z greater or equal than 1.50, 1.75 and 2 respectively and for each threshold we then selected the proteins that were unique for each fragment for each threshold. *omiXscore* calculations of ADAR and ADARB1 are interactions are respectively at http://crg-webservice.s3.amazonaws.com/submissions/2020-04/263420/output/index.html?unlock=f9375fdbf9 and http://crg-webservice.s3.amazonaws.com/submissions/2020-04/263140/output/index.html?unlock=bb28d715ea.

### GO terms analysis

*clever*GO ^49^, an algorithm for the analysis of Gene Ontology annotations, was used to determine which fragments present enrichment in GO terms related to viral processes. Analysis of functional annotations was performed in parallel with *GeneMania* ^57^. The link to *clever*GO analyses for fragment 1 is at http://www.tartaglialab.com/GO_analyser/render_GO_universal/3073/0d66e887c3/ (Z≥2).

### RNA and protein alignments

We used *Clustal W* ^39^ for 62 SARS-CoV-2 strains alignments and *T-Coffee* ^43^ for spike S proteins alignments. The variability in the spike S region was measured by computing Shannon entropy on translated RNA sequences. The Shannon entropy is computed as follows:

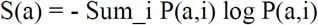

Where *a* correspond to the amino acid at the position *i* and P(a,i) is the frequency of a certain amino-acid *a* at position *i* of the sequence. Low entropy indicates poorly variability: if P(a,x) = 1 for one *a* and 0 for the rest, then S(x) =0. By contrast, if the frequencies of all amino acids are equally distributed, the entropy reaches its maximum possible value.

### Predictions of phase separation

*cat*GRANULE ^76^ was employed to identify proteins assembling into biological condensates. Scores > 0 indicate that a protein is prone to phase separate. Structural disorder, nucleic acid binding propensity and amino acid patterns such as arginine-glycine and phenylalanine-glycine are key features combined in this computational approach ^76^.

## SUPPLEMENTARY MATERIAL

**Supp. Figure 1**. We employed CROSSalign ^13,12^ was to compare the Wuhan strain MN908947 with other coronaviruses (>2000 strains, including SARS-CoV, MERS-CoV and coronaviruses having as host human or other species, such as bats). The result highlights that one of the most conserved region falls inside the Spike S genomic locus at nucleotides 22500-23000.

**Supp. Figure 2**. CROSS performances on betacoronavirus **(A**) 5’ end; (**B**) 3’ end ^33–36^. Using the confidence score corresponding to 1% of **Fig 1D**., we show that CROSS is able to identify double and single stranded regions with great predictive power (Area Under the ROC curve). We show here the performances for NC_006213 or Human coronavirus OC43 strain ATCC VR-759, NC_019843 or Middle East respiratory syndrome coronavirus, NC_026011 or Betacoronavirus HKU24 strain HKU24-R05005I, NC_001846 or Mouse hepatitis virus strain MHV-A59 C12 and NC_012936 or Rat coronavirus Parker.

**Supp. Figure 3**. Comparison between predicted and experimentally-validated interactions revealed by RAP-MS ^73^. From low (25%) to high (5%) confidence scores, the predictive power, measured as the Area Under the Curve (AUC) of Receiver Operating Characteristics (ROC), increases monotonically (HC corresponds to 10 nucleotides with highest/lowest scores), indicating strong agreement.

**Supp. Table 1**. Nucleotides 501-750 within fragment 23 contain structural motifs that recur in different coronaviruses. We hereby report the matches falling within Spike S region (+/-1000 nucleotides upstream or downstream) of annotated coronavirus, their NCBI codes, start and end coordinates as well as E-values.

**Supp. Table 2**. 1) catRAPID ^18,19^ score for interactions with fragment 1; 2) GO ^49^ and Uniprot annotations of viral proteins interacting with fragment 1 and; 3) catRAPID score for interactions with fragment 2; 4) GO annotations of viral proteins interacting with fragment 2; 5) catRAPID score for interactions with fragment 29; 6) GO annotations of viral proteins interacting with fragment 29;

**Supp. Table 3**. RBP interactions from Gordon et al. ^67^ classified according to catRAPID scores. GO ^49^ and Uniprot ^94^ annotations are reported.

**Supp. Table 4**. RBP interactions from Schmidt et al. ^73^. catRAPID scores and fold-changes are reported for different genomic regions (fragments ranging from 1 to 30).

**Supp. Table 5**. RBPs significantly enriched in the 5’ with GO ^49^ and Uniprot ^94^ annotations for HIV, HBV and Influenza A.

**Supp. Table 6**. Stress granule propensity computed with catGRANULE ^76^ for proteins with interactions score Z ≥ 1.5 (top-ranked) and Z<-1.5 (bottom-ranked). The two groups of proteins have the same number of elements.

