## Supplementary figures and images for "Structural analysis of SARS-CoV-2 genome and predictions of the human interactome"

### Supplementary Figure

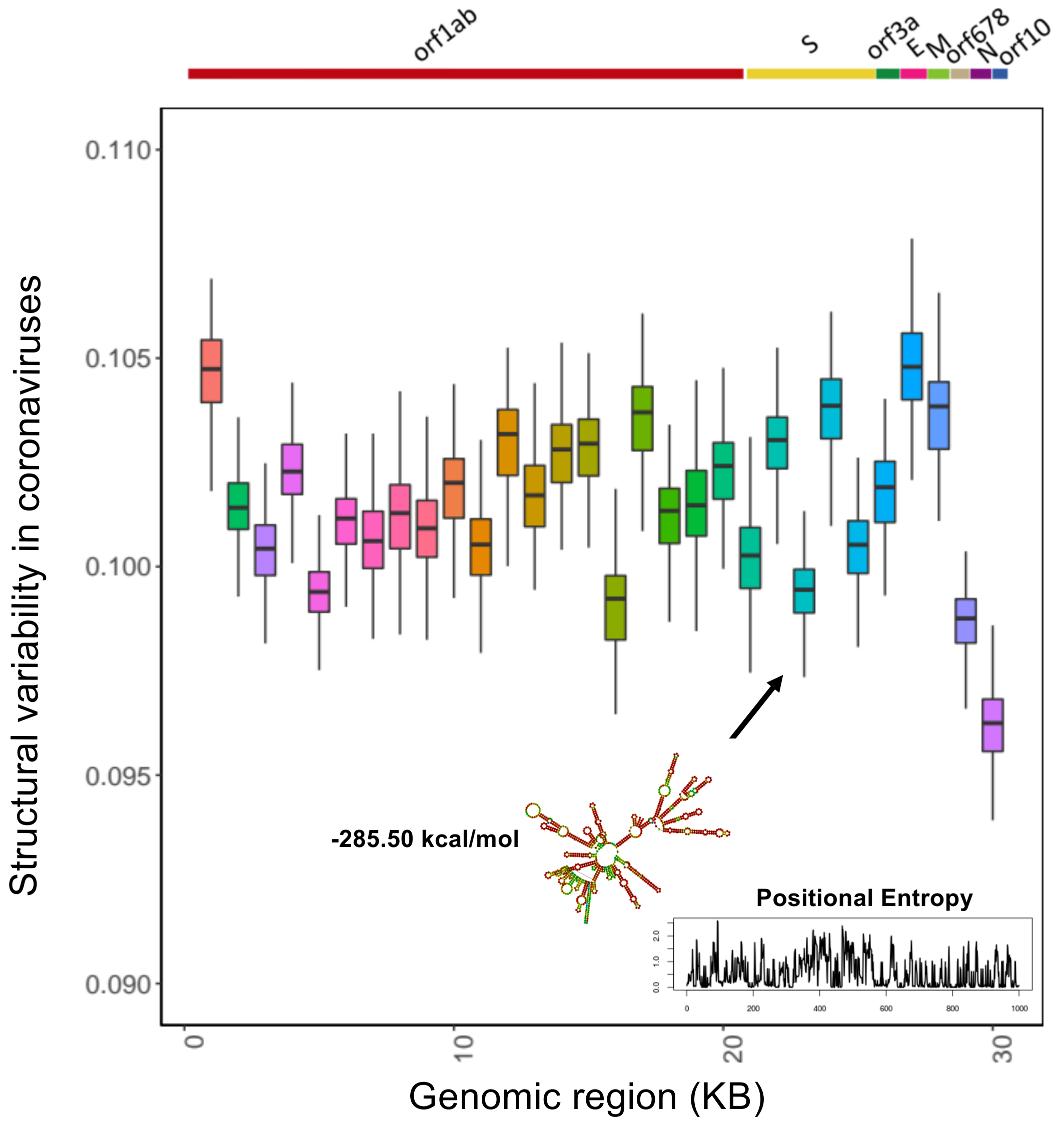

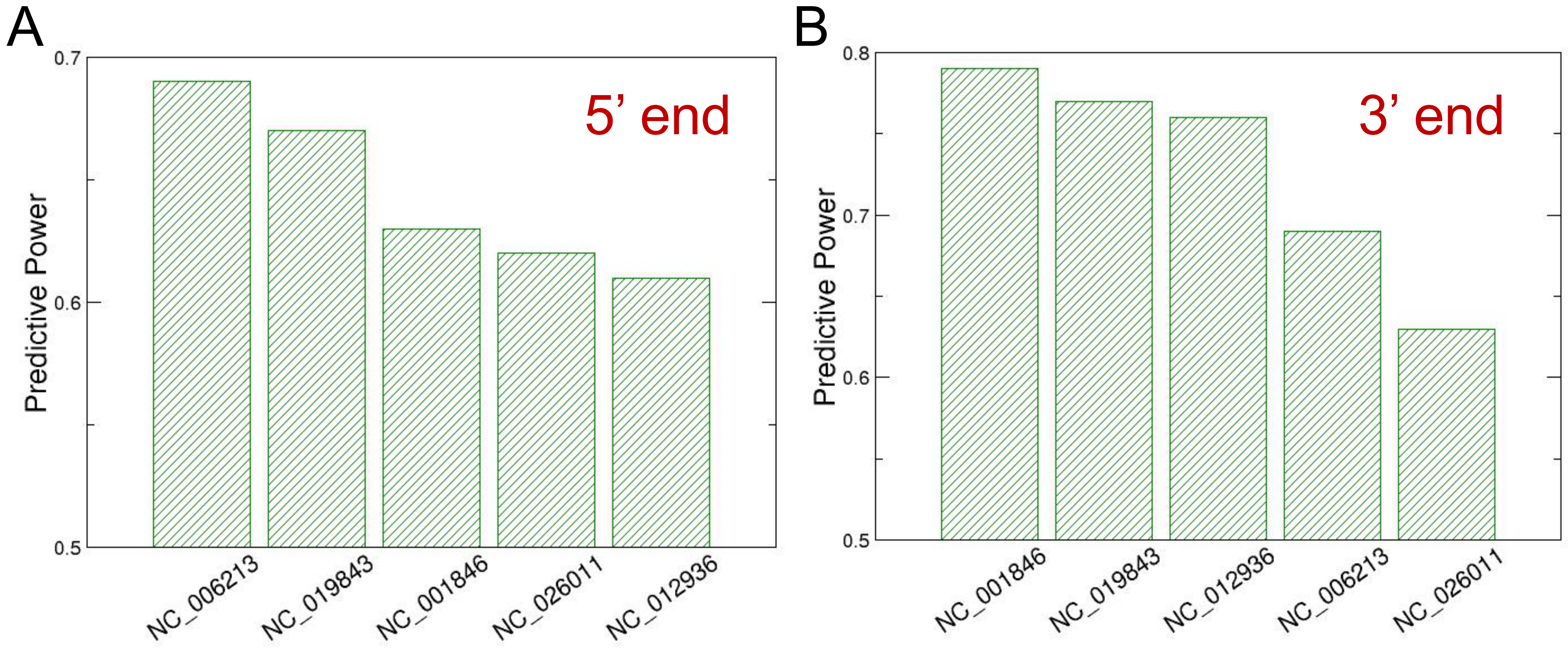

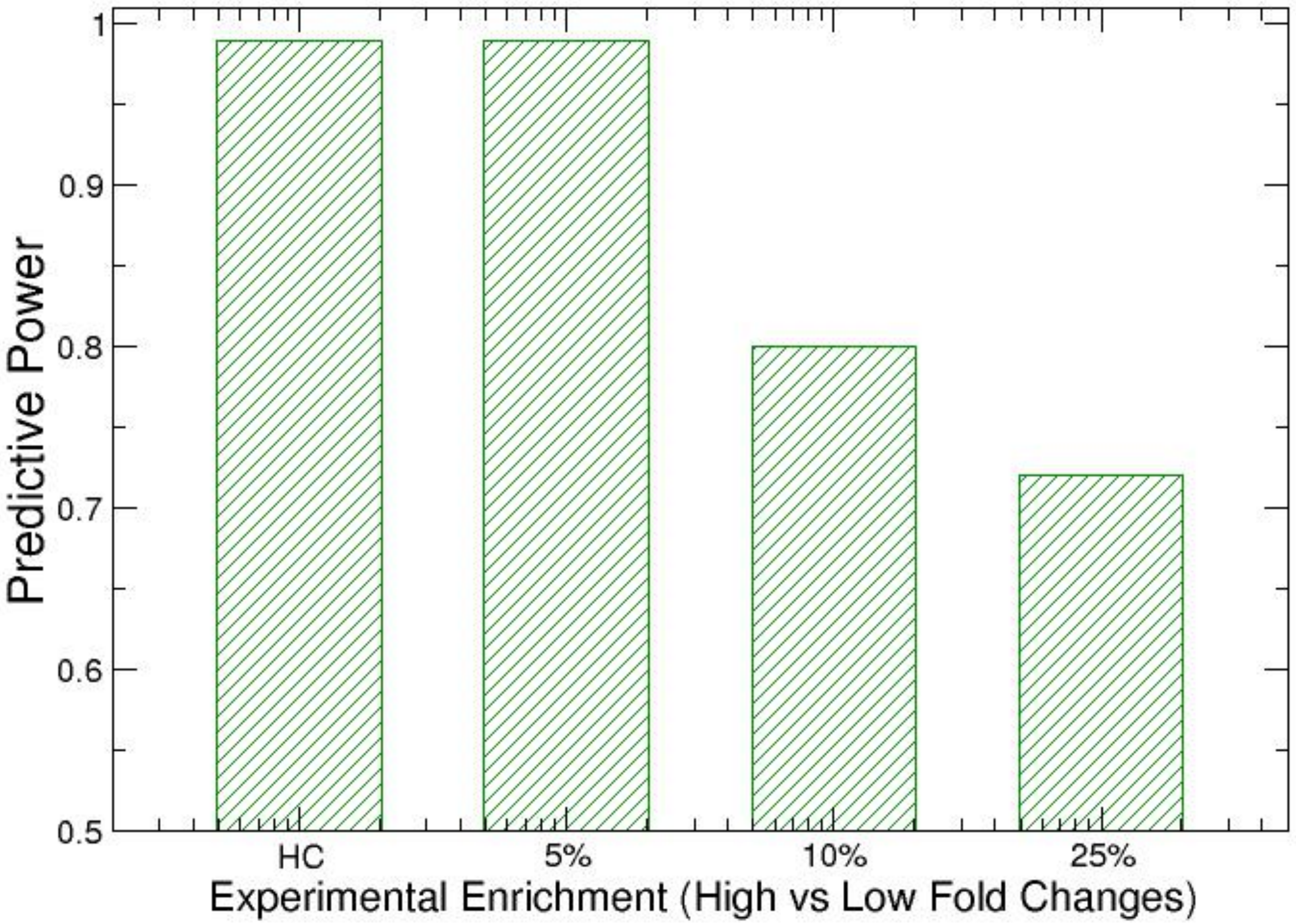
